# Perturbation of the titin/MURF1 signaling complex is associated with hypertrophic cardiomyopathy in a fish model and in human patients

**DOI:** 10.1101/680579

**Authors:** Yuta Higashikuse, Nishant Mittal, Takuro Arimura, Sung Han Yoon, Mayumi Oda, Hirokazu Enomoto, Ruri Kaneda, Fumiyuki Hattori, Takeshi Suzuki, Atsushi Kawakami, Alexander Gasch, Tetsushi Furukawa, Siegfried Labeit, Keiichi Fukuda, Akinori Kimura, Shinji Makino

**Affiliations:** Department of Cardiology, Keio University School of Medicine, 35-Shinanomachi Shinjuku-ku, Tokyo 160-8582, Japan; Division of Basic Biological Sciences, Faculty of Pharmacy, Keio University, Tokyo, Japan; Department of Molecular Pathogenesis, Medical Research Institute, Tokyo Medical and Dental University, Tokyo, Japan; Department of Interventional Cardiology, Cedars-Sinai Medical Center, 8700 Beverly Boulevard, AHSP A9229, Los Angeles, CA 90048, USA; Department of Biological Information, Tokyo Institute of Technology, Yokohama, Japan; Laboratory of Genome Diversity, Graduate School of Biomedical Science, Tokyo Medical and Dental University, Tokyo, Japan; Department of Integrative Pathophysiology, Medical Faculty Mannheim, Mannheim, Germany; Department of Bio-informational Pharmacology, Medical Research Institute, Tokyo Medical and Dental University, Japan; Keio University Health Centre, 35-Shinanomachi Shinjuku-ku, Tokyo 160-8582, Japan

**Keywords:** Genetics, Familial hypertrophic cardiomyopathy, Titin, Ubiquitin, Myocardial stiffness

## Abstract

Hypertrophic cardiomyopathy (HCM) is a hereditary disease characterized by cardiac hypertrophy with diastolic dysfunction. Gene mutations causing HCM have been found in about half of the patients, while the genetic etiology and pathogenesis remain unknown for many cases of HCM. To identify novel mechanisms underlying HCM pathogenesis, we generated a cardiovascular-mutant medaka fish non-spring heart (*nsh*), which showed diastolic dysfunction and hypertrophic myocardium. The *nsh* homozygotes had fewer myofibrils, disrupted sarcomeres and expressed pathologically stiffer titin isoforms. In addition, the *nsh* heterozygotes showed M-line disassembly that is similar to the pathological changes found in HCM. Positional cloning revealed a missense mutation in an immunoglobulin (Ig) domain located in the M-line-A-band transition zone of titin. Screening of mutations in 96 unrelated patients with familial HCM, who had no previously implicated mutations in known sarcomeric gene candidates, identified two mutations in Ig domains close to the M-line region of titin. *In vitro* studies revealed that the mutations found in both medaka fish and in familial HCM increased binding of titin to muscle-specific ring finger protein 1 (MURF1) and enhanced titin degradation by ubiquitination. These findings implicate an impaired interaction between titin and MURF1 as a novel mechanism underlying the pathogenesis of HCM.

## Introduction

Zebrafish is well established as a model vertebrate in gene function studies.^1, 2^ The medaka fish, *Oryzias latipes*, recently emerged as another important such model system, because of its smaller genome size and larger genome diversity.^3^ In particular, medaka is an excellent model for studying the cardiovascular system, because the embryos are transparent and able to tolerate an absence of blood flow under sufficient oxygen delivery by diffusion.^4^ It is therefore possible to attribute cardiac phenotype directly to particular genes by ruling out the secondary effect of hypoxia.

Hypertrophic cardiomyopathy (HCM) is characterized by cardiac hypertrophy accompanied by myofibrillar disarray and diastolic dysfunction. The first HCM-causing gene identified was a missense mutation in the cardiac β-myosin heavy chain gene.^5^ Such causative mutations were also identified in genes encoding contractile apparatus of sarcomere, such as α-tropomyosin, cardiac troponin T, and cardiac myosin-binding protein-C,^6^ whereby HCM was thought to result from compensation for dysregulated sarcomeric contractile function. An HCM-associated mutation^7-9^ identified in the titin gene (*TTN*) was the first causative gene not encoding contractile apparatus, which implied the existence of a novel pathogenesis except the compensation of systolic dysfunction.

Titin is the largest known single protein and, after myosin and actin, the third most abundant protein of vertebrate striated muscle. Titin connects the Z-disc to the M-line region where titin filaments extending from the opposite sides of half-sarcomeres overlap.^10, 11^ Titin provides a scaffold for the assembly of thick and thin filaments^12^ and passive stiffness during diastole.^13^ Recently, titin was proposed to function as a sarcomeric stretch sensor that regulates muscle protein expression and turnover.^14-16^ Titin carries a kinase domain at the M-line-A-band transition zone, and adjacent to the kinase domain is a binding site for the MURF1,^17^ where the mechanical load on the sarcomere is transmitted through titin and its interaction partners to the nucleus. Functional alteration due to the *TTN* mutations may increase passive tension upon stretching of the sarcomere.^8, 18^ Titin is extensively modular in structure and can be thought of as a microscopic necklace comprising up to 166 copies of 90-95 amino acid residue repeats from the immunoglobulin (Ig) domains.

Generally, the Ig domains located in the highly repetitive A-band region of titin share high structural similarity.^19^ However, a subset of Ig domains close to the titin kinase has unique structural features. For instance, Ig-A169 provides a binding site for MURF1,^17, 19^ a sarcomere-associated E3 ubiquitin ligase that conjugates ubiquitin to proteins destined for proteolysis.^20, 21^ MURF1 belongs to the MURF family of proteins, which are transcribed from three genes, form heterodimers,^17^ and interact with titin kinase. MURFs have also been proposed to regulate protein degradation and gene expression in muscle tissues.^15^

Here, we identified the *non-spring heart* (*nsh*) medaka fish, a cardiovascular mutant with a missense mutation in an Ig domain located at the M-line-A-band transition zone of titin. The *nsh* fish show diastolic dysfunction and hypertrophic heart. To decipher the roles of titin in these pathological effects, we focused on the M-line-A-band transition zone and identified two mutations in other Ig domains of titin in familial HCM patients. These titin mutations both increased the binding of titin to MURF1 and enhanced the degradation of titin by ubiquitination. This is the first report suggesting that the impaired binding of MURF1 to titin is a novel pathogenesis causing HCM.

## Results

### Medaka *nsh* mutants showed hypertrophic heart and disrupted sarcomeric structure

In a previous screening for ENU-induced mutations of medaka fish, we identified several mutants with abnormal heart development phenotypes.^25^ Among these, one mutant showed rigid hearts with hypertrophy. Because the mutant heart had lost elasticity and beat in a rigid manner, we designated it as *non-spring heart* (*nsh*). The *nsh* embryos were indistinguishable from their wild-type (WT) siblings up to the heart tube stage (48 hours) and there was no difference in the heart rate. However, after 72 hours, the *nsh* hearts showed increased thickening of the myocardial wall, especially in the atrium and even without blood flow (Figure 1A and 1B). Despite these abnormalities, the mutant embryos survived until 8 days post fertilization (dpf), a stage around hatching, in a relatively healthy condition without forming pericardial edemas.

**Figure 1.**
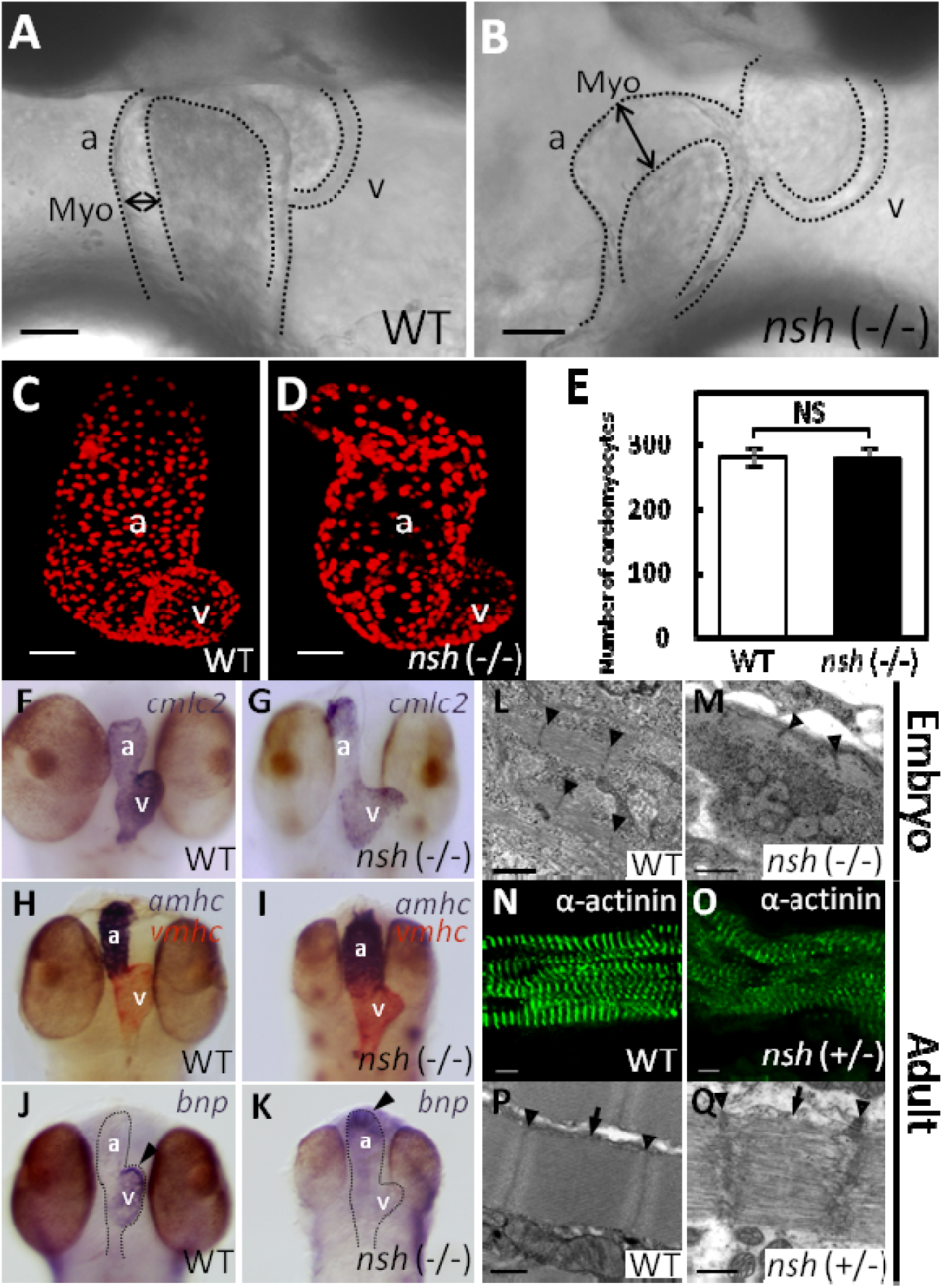
Hypertrophic myocardium and disrupted sarcomeric structure in the medaka *nsh* mutant heart. A and B, Morphology of the wild-type (WT) and *nsh* mutant heart at 3 days post fertilization (dpf). The mutant showed hypertrophic heart, especially in the atrium (double-headed arrow). The *nsh* heart lost elasticity and showed increased passive stiffness. Due to diastolic inability (see supplemental Movie 2), the blood flow was very slow or clogged around 3 dpf. C and D, Confocal Z-stacked images of the WT and *nsh* mutant hearts at 3 dpf, in which cardiomyocytes were visualized by the *Tg* signal (*cmlc2: DsRed2-nuc*). These reconstructed confocal images show approximately half of the total cardiomyocytes. E, Quantification of cardiomyocytes using the transgenic line. Cell counting was performed on the optical slices to attain an accurate count. There was no difference in the number of cardiomyocytes between the WT and *nsh* mutant hearts. NS, statistically not significant (n = 5 for WT and *nsh*). F through K, *In situ* RNA hybridization analysis showing cardiac myosin light chain 2 (*cmlc2*) (F and G), atrial and ventricular myosin heavy chains (*amhc* and *vmhc*, respectively) (H and I), in the WT and *nsh* mutant at 3 dpf. Normal sarcomeric protein expression was observed in the *nsh* mutant. J and K, Strong expression of brain natriuretic peptide (*bnp*) was detected at 3 dpf in the WT ventricle (J), whereas *bnp* expression was strong in the atrium of *nsh* mutants (K). L and M, Transmission electron microscopic findings of medaka embryonic cardiomyocytes at 5 dpf. Myofibrillar arrays were evident in the WT heart (L), but myofibrils were reduced with loosely arranged contractile filaments in the *nsh* mutant (*nsh* −/−) (M). Thick filament structures and integrity of M-line region were disrupted, and Z-discs were not parallel. N and O, Medaka adult cardiac myocytes in the WT and *nsh* heterozygotes (*nsh* +/−) were stained for α-actinin. Expression of sarcomeric α-actinin was maintained in the *nsh* heterozygous mutant, but fewer striations and patchy staining were observed (O). P and Q, Transmission electron microscopy showed the rupture of sarcomeric units, as well as irregular and widened Z-discs and M-lines in cardiomyocytes of the *nsh* heterozygotes. The integrity of the thick filaments is disrupted (Q). a, atrium; v, ventricle; Myo, myocardium. Z-discs (arrowhead) and M-lines (arrow) are indicated. Scale bars, 50 µm in (A through D), 1 µm in (L and M), 5 µm in (N and O), and 0.5 µm in (P and Q).

Blood flow was very slow or clogged around 3 dpf in the mutants due to defects in diastole (see supplemental Movies 1 and 2). However, hemoglobin activity was normal in both WT embryos and *nsh* mutants at 3 dpf based on an olto-dianisidine staining analysis (data not shown). After crossing with *fli1*-GFP transgenic medaka (Supplemental Figure I), alkaline phosphatase staining analysis revealed normal vascular endothelial cells at 3 dpf and normal blood vessel formation in the *nsh* mutants. To estimate the number of cardiomyocytes, we crossed the *nsh* mutants with a *Tg* medaka (*cmlc2:DsRed2-nuc*) expressing DsRed in the cardiomyocyte nuclei (Figure 1C and 1D). We counted the nuclei of cardiomyocytes using confocal images and found no difference in the numbers between WT and *nsh* mutants at 3 dpf (Figure 1C through 1E).

*In situ* hybridization with cardiac-differentiation markers, atrial and ventricular myosin heavy chains and cardiac myosin light chain, demonstrated normal expression of sarcomeric proteins in the *nsh* mutant (Figure 1F through 1I). A cardiac stretch marker, brain natriuretic peptide (*bnp*), is normally expressed at 3 dpf in the WT ventricle, but its expression was increased in the atrium of *nsh* mutants (Figure 1J and 1K).

We then examined the sarcomeric structures of embryonic cardiomyocytes by transmission electron microscopy. Sarcomeric units of the WT cardiomyocytes at 5 dpf showed highly organized, precisely aligned thick and thin myofilaments, flanked by Z-discs and M-lines (Figure 1L). In the *nsh* homozygotes, cardiomyocytes appeared abnormal with loosely arranged contractile filaments, where thick and thin filaments were disassembled and Z-discs were not parallel (Figure 1M). We next analyzed the sarcomeric structure of adult cardiomyocytes in the WT and *nsh* heterozygotes (Figure 1N and 1O respectively). Expression of sarcomeric α-actinin was maintained in the *nsh* heterozygotes, although with fewer striations and patchy staining (Figure 1O). Transmission electron microscopy demonstrated ruptured sarcomeric units with irregular and widened Z-discs and M-lines in the cardiomyocytes of *nsh* heterozygotes (Figure 1P and 1Q). We have analyzed body weight and heart weight ratio at 3∼4 month old adult medaka fish.and *nsh* heterozygotes heart showed heavier than WT heart (P=0.001, n=24).(Supplemental Figure II)

*nsh* did not show skeletal paralysis, however the *nsh* homozygous embryos had a slightly smaller body and skeletal myocytes demonstrated severely disturbed sarcomeric integrity with ruptured myofibrils. (Supplemental Figure III) Unlike in WT, staining for α-actinin was patchy in the *nsh* heterozygotes (Supplemental Figure III). By transmission electron microscopy, the skeletal myofibrils of *nsh* heterozygotes demonstrated occasionally disrupted M-lines, although they displayed well formed Z-discs and apparently mature myofibrils (Supplemental Figure III). Body size, movement, life span and fertility of the *nsh* heterozygotes were not affected (data not shown), although the abnormal sarcomeric structure was inherited as an autosomal dominant trait.

### *TTN* is the *nsh* gene

To identify the gene responsible for the *nsh* phenotype, we carried out positional cloning. We scored 2,016 fishes from *nsh* × *nsh*/+ crosses and found that the *nsh* phenotype was linked to the medaka linkage group 21, LG 21. Single-strand conformation polymorphism markers from introns 193-194 and 3’UTR of the *TTN* confined the *nsh* locus to a 19.0-kb critical region (Figure 2A). *TTN* was the only transcript identified in the *nsh* critical region. Analysis of cDNAs from the *nsh* mutant and several WT strains revealed an adenine to thymine transversion causing the substitution of D (aspartic acid)-23186 to V (valine) in the *nsh* titin (Figure 2B). Medaka titin is encoded by a single gene composed of 219 exons, and the *nsh* missense mutation was localized to *TTN* exon 204 near the MURF1 binding region that encoded an Ig domain (Figure 2C). Alignment of titin from several species revealed that the *nsh* mutation occurred at an evolutionary highly conserved amino acid residue (Figure 2D).

**Figure 2.**
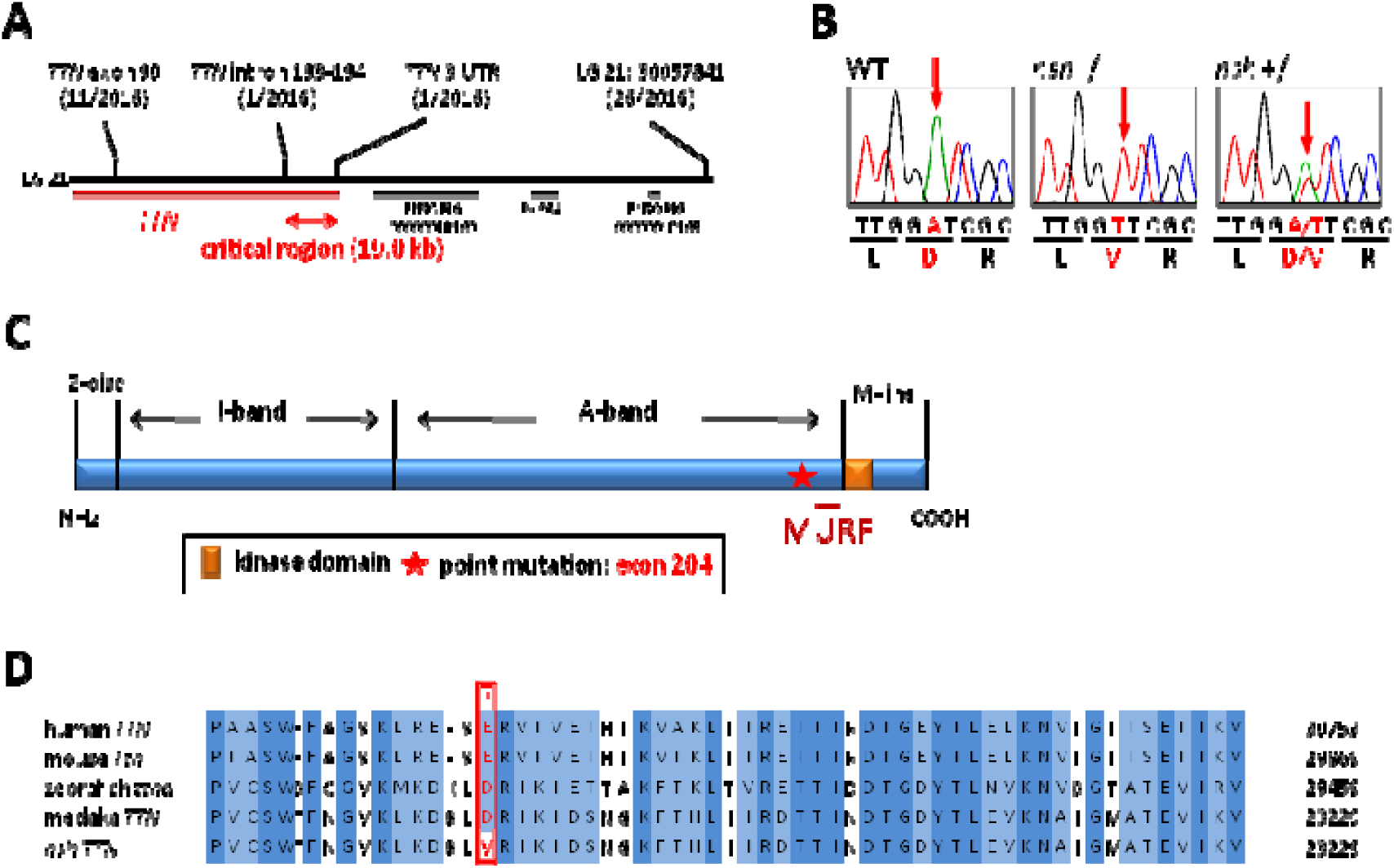
*TTN* is the *nsh* gene. A, Genetic mapping of the *nsh* locus on the medaka linkage group 21, LG 21. To define the *nsh* gene, we scored 2,016 fish from *nsh*/+ × *nsh*/+ mapping crosses. Single-strand conformation polymorphism markers from the intron 193-194 and 3’UTR of *TTN* confined *nsh* to a 19.0-kb critical region. B, DNA sequence analysis of cDNAs from the *nsh* and WT strains revealed an adenine to thymine mutation in *nsh*, causing D (aspartic acid)-23186 to V (valine) mutation (red arrow), located in exon 204. C, Modular structure of M-line-A-band transition zone of titin contains a binding site for muscle-specific ring finger protein 1 (MURF1). The missense mutation identified in *nsh* was located in an Ig domain near this site. D, Clustal W alignment of human, mouse, zebrafish, and medaka *TTN* sequences. Medaka titin and human titin proteins share 62% homology. GenBank or Ensembl accession numbers used for the analysis are as follows: human *TTN* (NM_133378), mouse *Ttn* (NP_035782), zebrafish *ttna* (DQ649453), and medaka *TTN* (ENSORLT00000022736). Alignment of titin protein from several species along with the *nsh* mutation showed that the mutation affected the evolutionary conserved amino acid residue (red asterisk). Identical and similar amino acids are indicated by dark blue and light blue boxes, respectively.

### Protein turnover of the mutated Ig domain titin at the M-line-A-band transition zone was faster than that of WT Ig domain titin

To investigate the molecular mechanisms of *TTN* mutation as the *nsh* gene, we performed RT-PCR analysis of the N-terminal *TTN*, A-I junction *TTN, TTN* exon 204, and titin kinase domain in WT and *nsh* mutant embryos at 3 dpf. This analysis revealed unchanged expression levels of *TTN* mRNA in the *nsh* mutant compared to WT (Figure 3A).

**Figure 3.**
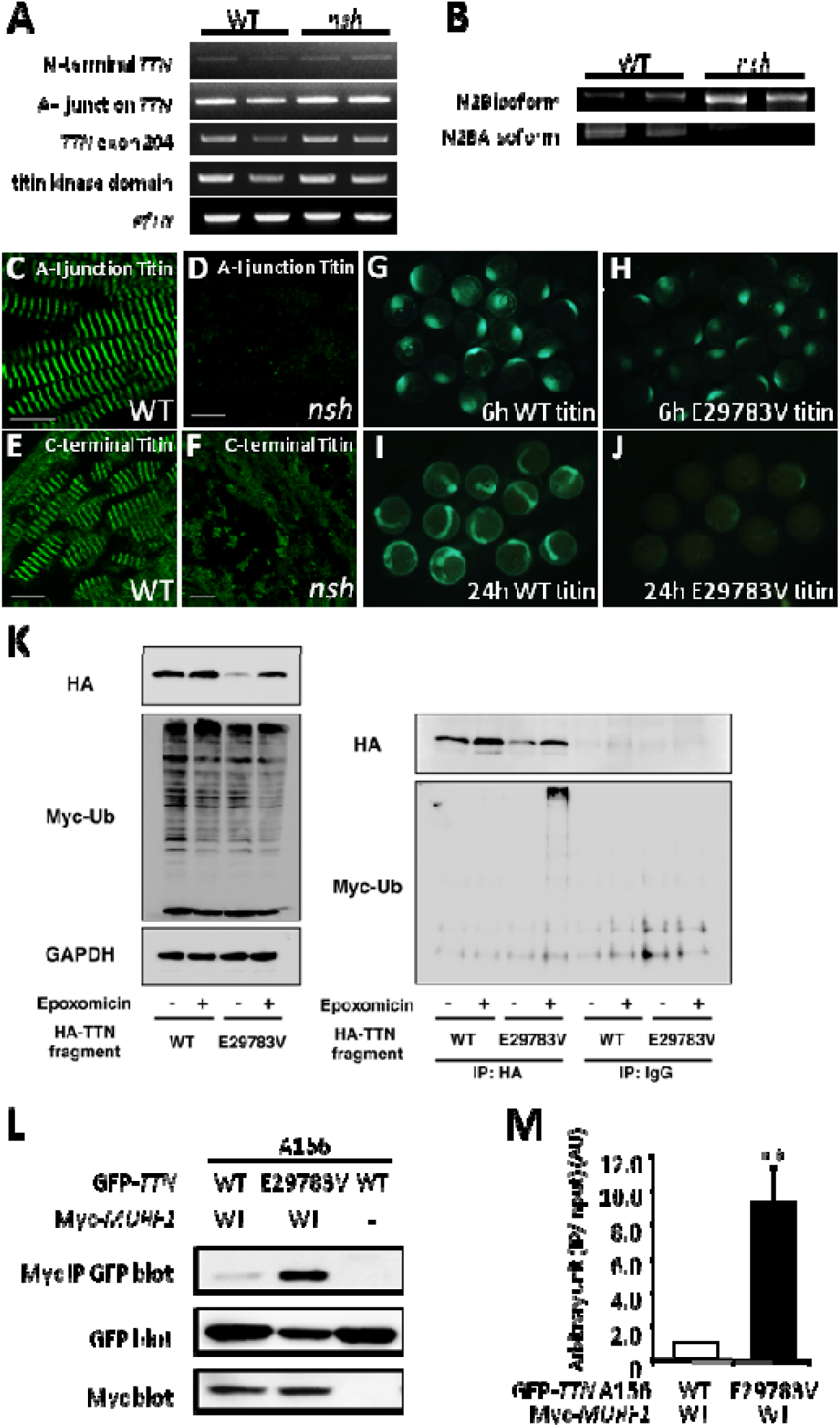
Mutant *TTN* Ig domain bound to MURF1 stronger and degraded faster *in vitro* and *in vivo*. A, RT-PCR analysis of *TTN* for N-terminal portion, A-I junction region, exon 204, and titin kinase domain in the WT and *nsh* mutant embryos at 3 dpf. Despite the presence of missense mutation, level of *TTN* mRNAs for these portions in the *nsh* mutant was similar to that in the WT. *ef1*α was used as a control. B, RT-PCR analysis of titin N2B isoforms in the WT and *nsh* mutants. N2B isoforms were increased. C through F, Embryonic skeletal muscle of the WT and *nsh* mutant at 3 dpf were stained with antibodies to A-I junction titin (C and D) and C-terminal titin (E and F). Despite the similar expression level of *TTN* mRNA, expression of titin containing the A-I junction isoform was reduced (D). Sarcomeric structures were disrupted in the *nsh* mutant (F). G through J, WT and mutant (E29783V which was the equivalent of D23186V mutation in medaka) Ig fragments mRNA injection to Medaka embryos. Despite the same amount of mRNA injection, mutant (E29783V) Ig fragments degraded faster than WT *TTN* Ig fragments. K, Western blots (WB) of HEK293FT cells transfected with HA-tagged WT and mutant (E29783V) Ig fragments with or without a proteosome inhibitor, epoxomicin. Myc-tagged ubiquitin was co-transfected to visualize the ubiquitination. WB of anti-HA IPs and control normal IgG IPs. Note the myc-labeled large molecules in mutant *TTN* transfectants were present only in the presence of epoxomicin. L, GFP-tagged *TTN*-A156 domain co-precipitated with myc-tagged MURF1 are shown (top panel). Expression levels of GFP-tagged *TTN*-A156 (middle panel) and myc-tagged MURF1 (lower panel) was confirmed by WB of whole cell lysates. Binding pairs were WT or E29783V mutant *TTN*-A156 in combination with MURF1. Dashes indicate no myc-tagged proteins (transfected only with pCMV-Tag3). M, Densitometric data obtained from the co-IP assays. Bar graphs indicate the amount of IPs normalized to the input amount of GFP-*TTN*. Value for WT *TTN*-A156 with MURF1 was arbitrarily defined as 1.00 arbitrary unit (AU). Data are represented as means +/- SEM. (n = 10). ** *P*<0.001 vs WT.

Titin isoform shifts have been reported in pathologic conditions.^34-38^ In cardiac muscle, the elastic region expresses two types of titin isoforms, termed N2B and N2BA, which arose from alternative splicing and result in titin molecules that contain either a unique N2B element and short PEVK region (N2B isoform) or unique N2B and N2A elements and a long PEVK region (N2BA isoform).^16^ Predominant expression of the N2B isoform results in stiffer cardiac tissue, while the expression of N2BA isoform results in more compliant cardiac myofibrils.^39^ Changes in the relative proportions of N2B and N2BA was paralleled by changes in diastolic stiffness.^40^ The N2BA/N2B ratio in myocardium under conditions of diastolic heart failure is lower than that observed during systolic heart failure.^41^ We detected different titin N2B isoform shifts between the WT and *nsh* mutants titin in the *nsh* heart, which would be associated with less elasticity and increased passive stiffness.

To analyze the expression of titin proteins in the *nsh* mutant, we conducted indirect immunofluorescence on 3 dpf embryos using an antibody to the A-I junction portion of titin. In WT, titin appeared as a regularly striated pattern between the sarcomeres of adjacent myofibrils (Figure 3C). However, the *nsh* mutants showed severely decreased titin immunofluorescence at the A-I junction (Figure 3D). Analysis of titin C-terminal region and α-actinin staining demonstrated an altered muscle organization, but there was no apparent decrease in expression of the titin C-terminal portion (Figure 3E, 3F and Supplemental Figures II). These findings indicated that certain titin isoforms at the A-I junction were selectively perturbed.

To further investigate the dominant-negative effect of mutant *TTN in vivo*, we overexpressed the Ig domain of *TTN* (GFP-tagged WT-*TTN*-A156 or E29783V-*TTN*-A156 construct, which was the equivalent of D23186V mutation in medaka) (Figure 3) by injecting mRNA corresponding to exon 204, in which the missense mutation was found, fused with the GFP fragment into one-cell-stage medaka embryos. Neither the WT-nor *nsh-*Ig domain induced a specific cardiac phenotypes (n = 200 in each case). However, in the *nsh*-Ig domain-injected medaka embryos, the GFP signal disappeared faster than in the WT Ig domain-injected embryos (Figure 3G through 3J), demonstrating that the *nsh*-Ig domain was susceptible to degradation and/or was unstable *in vivo*. The *nsh*-Ig domain fragments could not be observed 2 days after the injection, before the heart beating, and this could explain why we could not replicate the *nsh* phenotype in the injected embryos (Figure 3G through 3J).

Next we examined the faster turnover of the *nsh-*Ig titin domain, focusing on the ubiquitin proteasome (UPS) system because of the proximity of the *nsh* mutation to the MURF1 binding site. We found that the *nsh*-Ig domain protein was degraded faster via the UPS-dependent pathway *in vitro* than the WT Ig domain (Figure 3K), and that this effect was rescued by the UPS inhibitor, epoxomicin (Figure 3K), indicating the turnover of this specific titin Ig domain is controlled by the UPS. This regulation mechanism could involve MURF1 because it is thought to act in muscle as a rate-limiting E3 ligase to initiate UPS-mediated degradation.

### Medaka *nsh* mutants showed rigid hearts with normal heart rhythm

We quantified cardiac systolic and diastolic motion of WT and *nsh* mutants (see supplemental Movies 1 and 2) using high-speed video imaging and the analysis with motion vector prediction (MVP) method as described in the materials and methods section. This method could extract contraction and relaxation motions of cardiomyocytes non-invasively, and evaluate characteristics such as beating rate, orientation of contraction, beating homogeneity, and wave propagation of contraction.^42^ As shown in Figure 4B and C, maximum systolic and diastolic velocities of atrium were decreased from ∼183 μm/sec in WT embryos to ∼64 μm/sec in *nsh* mutant and from ∼140 μm/sec in WT embryos to ∼35 μm/sec in *nsh* mutant, respectively, while the heart rate between WT and *nsh* mutant was not changed significantly. Although the maximum systolic and diastolic velocities of WT ventricle showed similar values with those of atrium, such velocities of *nsh* mutant was lower than those of atrium, *i.e.*, ∼29 μm/sec for systole and ∼22 μm/sec for diastole (Figure 4C). These data demonstrated that *nsh* mutated gene was not required for the establishment of cardiac rhythm, but rather for control the cardiac elasticity (Figure 4B). To confirm if exaggerated degradation was responsible for the phenotype of *nsh*, we performed a rescue experiment with morpholino knockdown of MURF1. After the injection of morphlino we have analysis of cardiac movement by motion vector analysis. We could not suppress the *nsh* phonotype of cardiac hypertrophy even at a sub-lethal concentration however we could suppress the cardiac rigidity by MURF1 knockdown in *nsh* mutant. Cardiac hypertrophy is probably the later stages phenotype than cardiac rigidity and morphlino last in medaka embryo.

**Figure 4.**
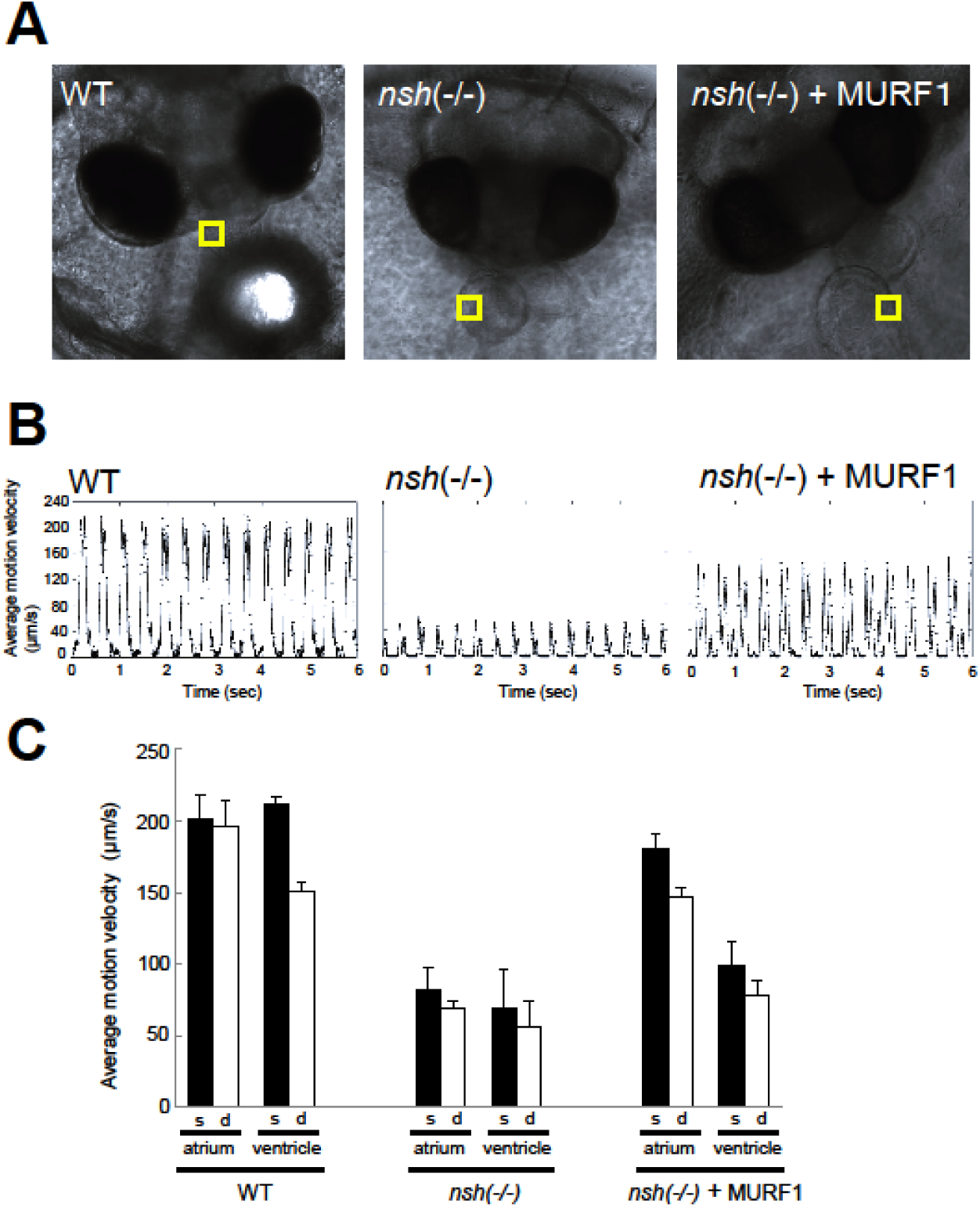
Medaka *nsh* mutants showed rigid hearts with normal heart rhythm. The beating motions of medaka atrium and ventricle were captured with high-speed camera and were analyzed with motion vector prediction (MVP) method. A, phase-contrast images of medaka embryos of WT (left), *nsh* mutant (middle) and *nsh* mutant + MURF1 Morpholino (right). The yellow squares in the images indicate the regions of interest (ROI) employed for the motion vector calculation of atrium wall. In B, typical beating profile of atrium observed for WT (left) *nsh* mutant (middle) and *nsh* mutant + MURF1 Morpholino (right). Motion velocity, the vertical scale, represents averaged motion vector length in the ROI. In both figures, each large peak followed by a small peak represents a single beat of atrium. Since we obtained the vector lengths simply from the square root of *x*^2^ + *y*^2^ (*x* and *y* represent the vector components), the systolic and diastolic peaks both had positive values. C, maximum velocities observed for systolic and diastolic motions in WT, *nsh* mutant and *nsh* mutant + MURF1 Morpholino were shown for atrium and ventricle. Systolic and diastolic velocities were decreased in *nsh* mutant and were partially rescued by MURF1 Morpholino, while the heart rate was not changed. Data are represented as means +/- SEM. (n = 3).

**Figure 5.**
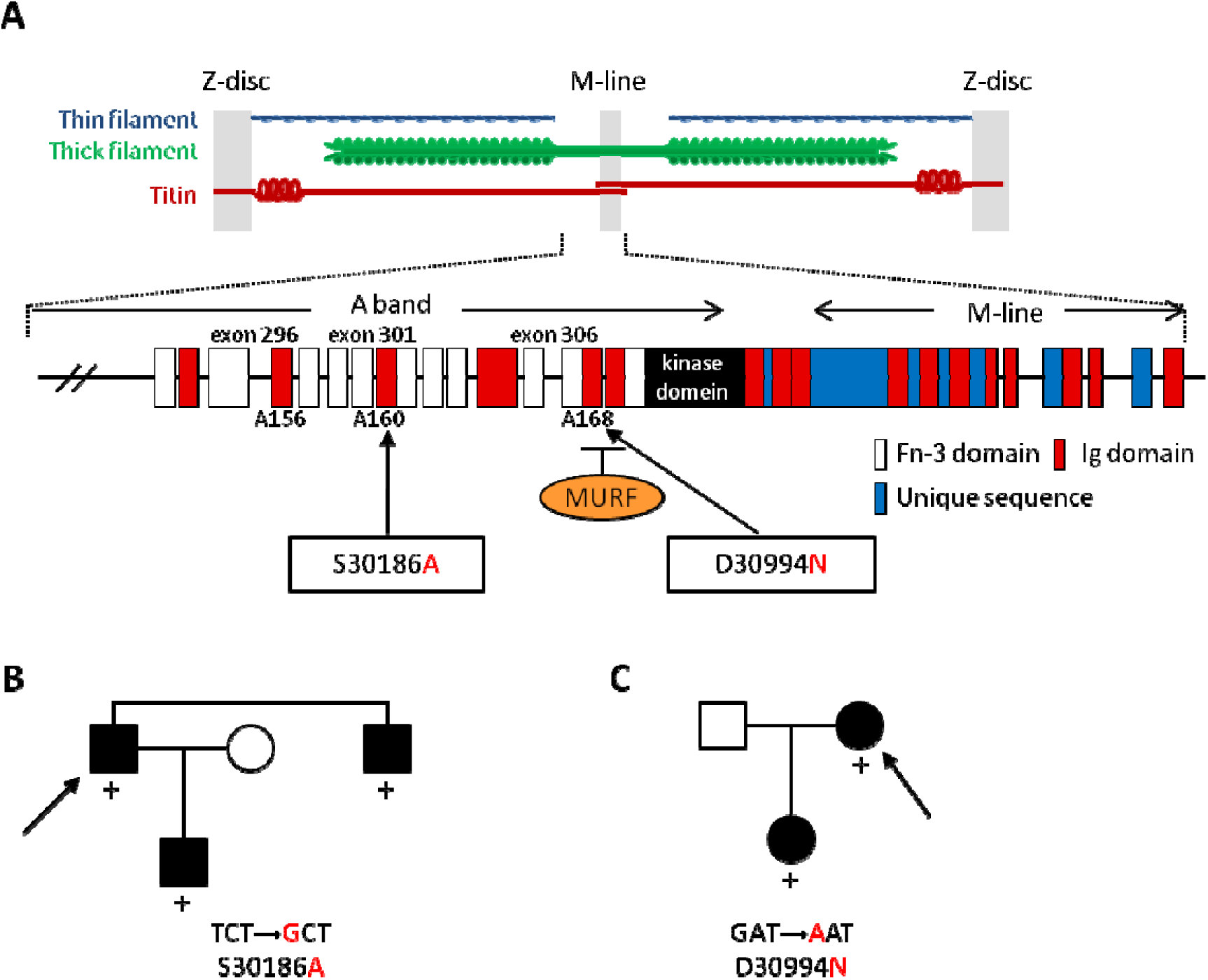
Mutations in the Ig domains of titin were associated with HCM. A, Schematic structure of titin at the C-terminal A-band and M-line region. C-terminal ends of titin meet in the M-line. Arrows indicate the position of two mutations found in familial HCM. The *nsh* missense mutation corresponds to exon 297 of human titin. Fn-3 indicates fibronectin type 3. B and C, Pedigrees of Japanese families with HCM; affected and unaffected members are indicated by solid and open symbols, respectively. Upon screening for mutations in exons 296-307 of *TTN*, which encode for the A-band region titin including the titin-kinase domain, two disease-associated mutations, S30186A (B) and D30994N (C), were identified. The latter mutation is located at the edge of the M-line region of titin and within a binding site for MURF1. Both mutations were found in the Ig domains of titin as indicated by red boxes in (A). The proband patient of each family is indicated by the arrow. Presence (+) of the mutations is noted.

Next we investigated the binding between MURF1 and Ig domain with or without the mutations using the coimmunoprecipitations (co-IP) assays, and found that co-IPs from cells transfected with the myc-tagged MURF1 construct in combination with GFP-tagged WT-*TTN*-A156 or E29783V-*TTN*-A156 construct revealed that the A156 domain bound MURF1 and that the E29783V mutation significantly increased the binding (9.23+/-2.11 AU, *P*<0.001) (Figure 4L and 4M). Notably, the expression level of mutant-*TTN*-A156 was lower than that of WT-*TTN*-A156 (Figure 6L).

**Figure 6.**
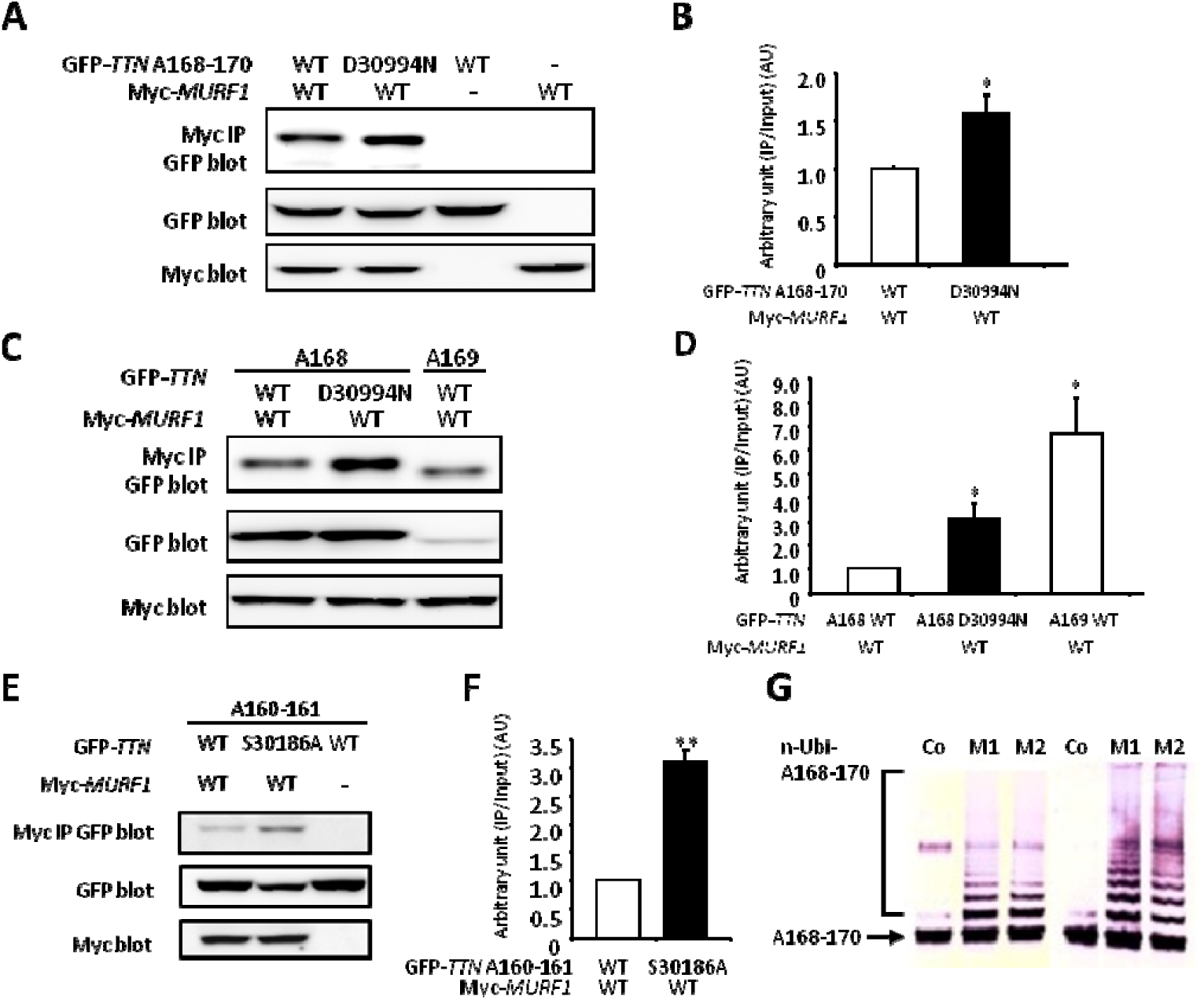
Altered binding of titin and MURF1 caused by the TTN mutations. A, GFP-tagged TTN-A168-170 domain co-immunoprecipitated with myc-tagged MURF1 is shown (top panel). Expression levels of GFP-tagged TTN-A168-170 (middle panel) and myc-tagged MURF1 (lower panel) were evaluated by WB of whole cell lysates. Binding pairs were WT or D30994N mutant TTN-A168-170 in combination with MURF1. Dashes indicate no GFP-tagged or myc-tagged proteins (transfected only with pEGFP-C1 or pCMV-Tag3, respectively). B, Densitometric data obtained from the co-IP assays. Bar graphs indicate the amount of precipitate normalized to the input amount of GFP-TTN. Value for WT TTN-A168-170 with MURF1 was arbitrarily defined as 1.00 AU. C, GFP-tagged TTN-A168 or TTN-A169 domains co-immunoprecipitated with myc-tagged MURF1 are shown (top panel). Expression levels of GFP-tagged TTN-A168 or TTN-A169 (middle panel) and myc-tagged MURF1 (lower panel) were evaluated by WB of whole cell lysates. Binding pairs were WT or D30994N mutant TTN-A168 in combination with MURF1, and WT TTN-A169 in combination with MURF1. D, Densitometric data obtained from the co-IP assays. Bar graphs indicate the amount of precipitate normalized to the input amount of GFP-TTN. Value for WT TTN-A168 with MURF1 was arbitrarily defined as 1.00 AU. Data are represented as means +/- SEM. (n = 10). * P<0.01 vs WT. E, GFP-tagged TTN-A160-161 domains co-precipitated with myc-tagged MURF1 are shown (top panel). Expression levels of GFP-tagged TTN-A160-161 (middle panel) and myc-tagged MURF1 (lower panel) was confirmed by WB of whole cell lysates. Binding pairs were WT or S30186A mutant TTN-A160-161 in combination with MURF1. Dashes indicate no myc-tagged proteins (transfected only with pCMV-Tag3). F, Densitometric data obtained from the co-IP assays. Bar graphs indicate the amount of IPs normalized to the input amount of GFP-TTN. Value for WT TTN-A160-161 (F) with MURF1 was arbitrarily defined as 1.00 AU, respectively. Data are represented as means +/- SEM. (n = 10). * P<0.01 vs WT and ** P<0.001 vs WT. G, Titin A168-170 was ubiquitinated by MURF1 and MURF2. The titin fragment A168-170 containing the MURF1 binding site was used as a substrate for *in vitro* ubiqutination assays. Addition of recombinant MURF1 or MURF2 catalyzes multi-ubiquitination of titin fragments. Lanes: Co, control: no E3 ligase added; M1: inclusion of MURF1; M2: inclusion of MURF2. After incubation, A168-170 fragments were detected by WB, using specific antibodies to titin A169 (left), and A168-170 (right).

Taken together, our data indicate that the Ig domain of the M-line-A-band transition zone of titin plays an important role in maintaining and organizing the myofibril structure. The severely reduced titin protein levels in combination with the genetic linkage data strongly implicated the *TTN* mutation as causative for the *nsh* phenotype. The alternations in N2B/N2BA titin isoforms in the *nsh* phenotype in turn underscored the probable key role of titin in regulating and maintaining cardiac muscle elasticity.

### Mutational analysis of the human titin gene around the MURF1 binding domain in patients with familial hypertrophic cardiomyopathy

The proximity of the *nsh* mutation to the titin-MURF1 binding domain and the cardiac phenotype in mutant medaka implicated the titin M-line-A-band transition zone as a candidate region for mutations associated with HCM in humans. We therefore analyzed a total of 96 genetically unrelated Japanese patients with familial HCM, who carried apparent family history of HCM consistent with an autosomal dominant trait. They were diagnosed with HCM according to the diagnostic criteria as described previously.^33^ They had been analyzed for mutations in the known disease genes for HCM, i.e., genes for contractile elements (MYH7, TNNT2, MYBPC3, TNNI3, MYL2, MYL3, ACTC1, TNNC1), Z-disc elements, and I band elements, and no disease-associated mutation was found.^33^ Control subjects were 400 healthy Japanese individuals selected at random.

Exon 204 of medaka *TTN* containing the *nsh* mutation corresponds to exon 297 of human *TTN*, thus we searched for mutations in the patients by direct sequencing of exons 296-307 of *TTN* (GenBank Accession No. NM_133378), which encoded for the A-band-M-line transition region of titin including the titin-kinase domain. We identified two disease-associated missense mutations (Figure 5A): TCT→GCT leading to the replacement of serine with alanine at position 30186 (S30186A) in exon 301 (Figure 5B), and GAT→AAT with the exchange of aspartic acid for asparagine at position 30994 (D30994N) in exon 306 (Figure 5C). The proband patients and their affected relatives were heterozygous for the mutations, and both mutations affected highly conserved amino acids. Quite interestingly, the latter mutation in the M-line-A-band transition zone of titin was mapped within the binding site for MURF1. It should be noted here that all three mutations found in medaka *nsh* mutant and human HCM patients were located in the Ig domains (Figure 5A).

### Altered binding of titin to MURF1 caused by the *TTN* mutations

Based on the D30994N mutation being located in the MURF1 binding Ig domain of titin, A168-170, we investigated a possible functional change in binding to MURF1 due to the mutation. GFP-tagged WT- or D30994N-*TTN*-A168-170 construct was co-transfected with a myc-tagged MURF1 construct into COS-7 cells. Western blot (WB) analysis of co-IPs of the transfected cells using anti-myc antibody demonstrated that the D30994N mutation significantly increased binding to MURF1 (1.58+/-0.19 AU, *P*<0.01) (Figure 6A and 6B). The A168-170 domain contains two Ig domains, A168 and A169, of which the A169 domain is the predominant binder of MURF1,^19^ despite the D30994N mutation being located in the A168 domain. To further characterize the increased binding of titin to MURF1 in the presence of the D30994N mutation, we performed additional WB analyses of co-IPs from the cells transfected with the myc-tagged MURF1 construct in combination with GFP-tagged WT-*TTN*-A168, D30994N-*TTN*-A168, or WT-*TTN*-A169 constructs. WT-*TTN*-A169 showed significantly higher binding to MURF1 than WT-*TTN*-A168 (6.69+/-1.53 AU, *P*<0.01) (Figure 6C and 6D). On the other hand, the D30994N mutation also significantly increased the binding of A168 domain to MURF1 (3.06+/-0.68 AU, *P*<0.01) (Figure 6C and 6D).

Because the other mutation also located in the Ig domain, we also investigated the binding between MURF1 and other Ig domain with or without the mutation using the co-IP assays. The co-IPs from cells transfected with the myc-tagged MURF1 construct in combination with GFP-tagged WT- or S30186A-*TTN*-A160-161 construct revealed that the A160-161 domains also bound MURF1 and that the S30186A mutation significantly increased the binding (3.11+/-0.71 AU, *P*<0.001) (Figure 6E and 6F). Notably, the expression level of mutant-*TTN*-A160-161 was lower than that of WT-*TTN*-A160-161 (Figure 6E). These data suggested that mutations in the Ig domains of M-line-A-band transition zone enhanced the interactions with MURF1. This raises an interesting possibility that the association of MURFs with the M-line regions could link ubiquitin-transfer pathways to the titin filament system, which may relate to the physiological turnover of titin in myofibrils. To test this hypothesis, we performed an *in vitro* ubiquitination assay of titin fragment. The titin fragment A168-170 containing the MURF1 binding site was used as a substrate. Addition of recombinant MURF1 or MURF2 indeed catalyzed multi-ubiquitination of titin fragment (Figure 6G). Moreover to confirm this idea, we also performed immuno-electron microscopy with anti-ubiquitin antibodies in WT and MURF1 knock-out mouse.^20^ Ubiquitin signals was detected at the Z-line, central I-band and M-line regions in WT sarcomere. It was hard to quantify the signals, however ubiquitin seemed to be reduced in MURF1 knock-out mice at the M-line region (data not shown). Endogenous MURF1 levels were tightly regulated,^17^ in combined with ubiquitination of titin A168-170 fragment by MURFs (Figure 6G), reduced ubiquitin signals of immuno-electron microscopy and increased binding of titin and MURF1 (Figure 4L, 6A, 6C and 6E) suggested that the interactions of MURF1 with titin are critical for maintaining the stability of titin.

## Discussion

We generated a cardiovascular-mutant *nsh* medaka fish by ENU mutagenesis that developed hypertrophy and diastolic dysfunction, and identified a causative mutation in a region of titin around the MURF1 binding site. The *nsh* homozygous fish showed loosely arranged contractile units, and died around 8 dpf. The *nsh* heterozygotes showed normal life span and fertility, but they exhibited M-line disassembly in myofibrils, indicating that the phenotype of M-line disassembly was inherited as a dominant trait.

Due to the extremely large size and presence of multiple isoforms, it has been difficult to study the structure and function of titin. A knock-in mouse of eGFP to titin gene showed that sarcomeric titin is more dynamic than previously suggested.^42^ The large cardiac titin N2BA isoform is rapidly replaced by the smaller N2B isoform both after birth and with re-expression of the fetal gene program in cardiac pathologic conditions.^34-38^ The isoform switch plays an important role in adjusting diastolic function. It has been used with knock-out technology to gain insight into the roles of titin in sarcomere assembly, stability and mechanotransduction during muscle pathophysiology.^43-45^ Based on these studies, it was proposed that titin provides a scaffold for the developing sarcomere,^12^ and that passive forces maintain the correct localization of M-lines and tether various signaling and structural molecules.^11^ Multiple interacting proteins integrate titin into the sarcomere and modulate the titin filament with links to basic cellular functions including trophic signaling and energy balance.

Herman et al.^46^ recently showed by next-generation sequencing that *TTN* truncating mutations are the most common known genetic cause of dilated cardiomyopathy (DCM). Their findings revealed that mutations associated with DCM were overrepresented in the titin A-band, but were absent around the M-band regions of titin. One may therefore speculate that some truncated titin peptides are degraded via the RNA- and protein-surveillance pathways in DCM. The consequently decreased titin levels could limit sarcomere formation resulting in cardiac dysfunction. However, this is not the case for previously reported *TTN* mutations that delete only part of the M-band portion of titin^47^; immunohistochemical studies showed that some of these carboxy-terminal truncated titin proteins are integrated into the sarcomere. If more proximal *TTN* truncating mutations caused DCM through haploinsufficiency, the distribution of such mutations would be rather uniform across the susceptible portion of titin.

The current study of our HCM families found point mutations around the M-line-A-band transition zone, which is implicated in sensing and modulating sarcomeric force. These mutations also induced high binding affinity between titin and MURF1. Using the medaka fish and our *in vitro* system, we showed that mutated titin fragments are degraded faster by ubiquitination. This study indicate that titin proteins found in families with HCM, as in the carboxy-terminal titin truncations studied, are integrated into sarcomeres and cause HCM via a dominant negative mechanism. Our results indicated that titin protein mutations in HCM families could be incorporated into the sarcomere and impair MURF1 binding, resulting in HCM through disturbance of the normal turnover of titin and abnormal sarcomere stiffness.

Until now, altered binding of MURF1 to titin has not been identified to be closely associated with human HCM or diastolic dysfunction. In this study, we analyzed exons 296-307 (M-line-A band transition zone) of *TTN* for mutations in patients with familial HCM and identified two disease-associated missense mutations in the Ig domains. One of the *TTN* mutations in familial HCM was in a binding site for MURF1. Although MURF1 binds to the Ig domain of the M-line-A-band transition region, A168-170, MURF1 was not considered to target titin for degradation.^17^ Data in this study and previous studies indicate MURF1 may regulate the turnover of critical sarcomeric proteins^21^ including some titin isoforms (Figure 4C and 4D).^20, 32^ We also found that all the *TTN* mutations associated with HCM both in medaka and human in the Ig domains of the M-line-A-band transition zone increased the binding of MURF1 to titin (Figure 6A, 6C and 6E). In addition, the *TTN* constructs carrying the HCM-associated mutations expressed less amount of protein when they were injected into medaka embryos and transfected into cells (Figure 4G, 4H, 4I, 4J, 4L and 6). Although the molecular mechanisms underlying this reduced expression remain to be clarified, it seems to be caused by accelerated degradation by the ubiquitin proteasome pathway (Figure 4K and 6G). Further, the knockdown of MURF1 in *nsh* mutants partially recovered the cardiac systolic and diastolic velocity. These studies therefore suggests the inhibition of MURF1 function as a therapeutic intervention for HCM patients.

In summary, we have identified *TTN* mutations that increased the binding of titin to MURF1 in a medaka model and in patients with HCM. There are several pathways underlying the pathogenesis of HCM, including abnormal power generation, power transmission, calcium sensitivity, energy homeostasis and stretch response. MURF1 may be involved with a novel pathway responsible for both the regulation of myofibril turnover and isoform shift as well as muscle gene expression, because MURF1 exists in multiple locations within the cell including the cytosol, sarcomere, and nucleus. Our observations suggested that pathogenic mechanisms underlying the titin M-line-A-band transition zone-related HCM cases might be a consequence of abnormal protein turnover and impaired adjusting diastolic function regulated by titin and MURF1.

## Materials and methods

### Fish strains and maintenance

Medaka fish (*Oryzias latipes*) were maintained at 28.5°C in a recirculating system with a 14-h/day and 10-h/night cycle. The Qurt strain, derived from the southern Japanese population was used for the genetic screen. Another strain, Kaga, which came from the northern population in Japan, was used for the mapping cross. The medaka heart mutant *non-spring heart* (*nsh*) was isolated by mutational screening.^22^ Fertilized eggs were collected and then incubated at 28.5°C in medaka Ringer’s solution (0.65% NaCl, 0.04% KCl, 0.011% CaCl_2_, 0.01% MgSO_4_, 0.01% NaHCO_3_, and 0.0001% methylene blue). Developmental stages were determined by their morphology according to the medaka stage table described by Iwamatsu (2004).^23^ The fli1-GFP transgenic line was kindly gifted by Dr. A. Kudo.^24^ All experimental procedures and protocols were approved by the Animal Care and Use Committee of Keio University and conformed to the NIH Guidelines for the Care and Use of Laboratory Animals.

### Counting cardiomyocytes *in vivo*

The *DsRed2-nuc* gene under control of the zebrafish *cmlc2* promoter transgenic strain^25^ was used for scoring cardiomyocyte numbers. The wild-type and *nsh* homozygous hearts carrying *Tg (cmlc: DsRed2-nuc)* were dissected, fixed with 4% paraformaldehyde (PFA) in phosphate-buffered saline (PBS) overnight at 4°C, and then mounted in 1% agarose gel. We routinely prepared 50-100 optical sections from one heart sample, and counted cells on the merged images from one in every 25 sections. Data were presented as mean +/- SEM. Student’s t-test was used to test statistical significance.

### Whole-mount RNA *in situ* hybridization (ISH) analysis

Whole-mount ISH analysis was performed as described previously.^26^ After staining, tissues were fixed with 4% PFA in PBS for color preservation, equilibrated with 80% glycerol, and then mounted on slide glass for microscopic observation. The following primers were used for cloning the probes by polymerase chain reaction (PCR): for cardiac myosin light chain2 (*cmlc2*); 5’-AATGTCTTTTCCATGTTTGAGC-3’ and 5’-CTCCTCTTTCTCATCCCCATG-3’, for atrium myosin heavy chain (*amhc*); 5’-TGCACTGATGGCTGAATTTG-3’ and 5’-ACTTGATCTACACCTTGGCC-3’, for ventricular myosin heavy chain (*vmhc*); 5’-GCTGAGATGTCCGTGTATGGTGC-3’ and 5’-GCTCCTCACGAGGCCTCTGCTTG-3’, and for brain natriuretic peptide (*bnp*); 5’-GATCCATCCATCCATCATCC-3’ and 5’-TGATACTTTAAAGACACAATGTCCAA-3’. The PCR products were cloned into standard vectors and used to prepare the digoxigenin (dig)-labeled RNA probes.

### Histology and electron microscopy

We fixed medaka embryos with 4% PFA in PBS overnight at 4°C, before dehydrating them in 70% ethanol, mounting in paraffin, and sectioning at 5 µm thickness. Sections were stained with hematoxylin and eosin (HE) and viewed using a BIOREVO optical microscope (BZ-9000; Keyence, Japan). For transmission electron microscopy, samples were prepared as described previously,^27^ sectioned using an RMC MT6000 ultramicrotome, and visualized using the JEM-1230 (JEOL) electron microscope.

### Immunohistochemistry

Medaka embryos were fixed in 4% PFA/PBS overnight at 4°C. Following fixation and removal of the egg chorion with forceps, samples were cryoprotected in sucrose at 4°C overnight, embedded in OCT compound (Sakura Finetec), and snap-frozen in hexane at −80°C. The sections were incubated in primary antibodies overnight at 4°C, washed three times for 5 min each in PBS, and then incubated in the appropriate secondary antibodies (Invitrogen) for 1 h at room temperature. Stained samples were observed by confocal laser-scanning microscopy (LSM510; Carl Zeiss, Jena, Germany). The following primary antibodies were used at the indicated dilutions: anti-α-actinin (Sigma; A7811) at 1:300, anti-A-I junction Titin [T11 (Sigma; T9030)] at 1:100, and anti-C-terminal Titin [Titin H-300 (Santa Cruz; sc-28536)] at 1:100.

### High-speed video microscopy

Medaka embryos were placed in a bottom of the well (24-multiwell plate) and incubated with 500μl of medaka Ringer’s solution. Embryos were arranged to direct their heart toward bottom of the plate so that the cross-sectional heart images for each embryo can be observed from similar direction. A high-speed digital CMOS camera (KP-FM400WCL, Hitachi Kokusai Electric, Tokyo, Japan) was mounted on an inverted microscope (Eclipse TE2000, Nikon, Tokyo, Japan), which was equipped with an *xy* scanning stage (Bios-T, Sigma Koki, Tokyo, Japan). The microscope was also equipped with a stage-top mini-incubator (WSKM, Tokai Hit, Shizuoka, Japan), which maintained the temperature of the culture plate at 28.5°C. Video of medaka embryos was recorded as sequential phase-contrast images with a 10× or 20× objective at a frame rate of 150 fps, a resolution of 2048×2048 pixels, and a depth of 8 bits.

### Evaluation of beating heart with MVP algorithm

To evaluate cardiac systolic and diastolic motion of medaka heart from the high-speed video imaging of the embryos, we have applied MVP method as previously reported.^28, 29^ Briefly, each frame was divided into square blocks of *N* × *N* pixels (Supplementary Figure IV). Then, for a maximum motion displacement of *w* pixels per frame, the current block of pixels was matched to the corresponding block at the same coordinates in the previous frame within a square window of width *N* + 2*w*. (Optimal values of *N* and *w* for the cardiac motion detection may vary with the observation magnification and resolution of the employed camera. Here, we set *N* = 16 and *w* = 4. These values were determined empirically based on the throughput speed of calculation and accuracy of the block-matching detection.) The best match on the basis of a matching criterion yielded the displacement. The mean absolute error (MAE) was used as the matching criterion. The matching function is given by

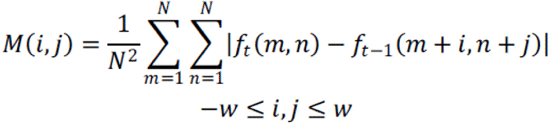

where *f*_*t*_(*m, n*) represents the intensity at coordinates (*m, n*) in the current block of *N* × *N* pixels and *f*_*t-1*_ (*m* + *i, n* + *j*) represents the intensity at new coordinates (*m* + *i, n* + *j*) in the corresponding block in the previous frame. We performed this calculation for every 4 × 4 pixels in the region of interest (ROI).

### Genetic mapping and positional cloning

Genetic mapping was performed as described previously.^30^ Briefly, an *nsh* mutant on the Qurt genetic background was crossed with the Kaga strain to obtain F1 offspring, which were then crossed to obtain the *nsh* mutant embryos for mapping. A pooled bulk segregation analysis was performed to localize the *nsh* mutation on the medaka linkage group.^30^ Further mapping was also done using custom-designed markers based on the sequence polymorphisms that differed between the Qurt and Kaga strains. The following primers and restriction enzymes were used for the mapping: for *TTN* exon 90; 5’-AACTTCACAGGCTTCAGGTC-3’ and R: 5’-CAGCATCTGTGACGTAGCTC-3’ (Hae III), for *TTN* intron 193-194; 5’-CCGTGTAACCAACCTCACTG-3’ and 5’-GGCTTGGTCCATTCAAGTGT-3’ (Hae III), for *TTN* 3’UTR; 5’-CAGGAATTGGTCCTGAAAGG-3’ and 5’-TCGACTTGGGTGTTTTCAGA-3’, for LG 21_30057841; 5’-AATAGGTTGAGGCTGCATGG-3’ and 5’-TTCACTGGTTCTTGGCGTTC-3’ (Hae III). The reference coding sequence of *TTN* cDNA was in the Ensembl DNA sequence database under the accession number ENSORLT00000022736.

### RT-PCR analysis

Hearts were harvested from wild-type and *nsh* mutant embryos (n = 50 for each) at 3 days post fertilization (dpf). Total RNA was extracted using Trizol reagent (Invitrogen). For RT-PCR analysis, we carried out oligo-dT primed reverse transcription on aliquots (1 µg) of total RNA and used 1/10 of the single-strand cDNA products for each PCR amplification. Gene-specific primers were designed based on published sequences. *ef1*α was used as an internal control. The following primers were used for RT-PCR analysis; for N-terminal *TTN*; 5’-GCTCTGCACAAAGAGCTGTC-3’and 5’-GCAAGATGTGCTGATGGTTC-3’, for A-I junction *TTN*; 5’-GCAAGCCAAGAACAAGTTTGA-3’ and 5’-AGAAAAGGTCGGGACAGAGG-3’, for *TTN* exon 204; 5’-TTGTTCATGTCCATCGTGGT-3’ and 5’-GCTCCTCTGGAGTTGATTGC-3’, for titin kinase domain; 5’-AATGTTGCAAGGCACAAAAA-3’ and 5’-AGCCTCAAGGCTAATGTCCT-3’, for *ef1*α; 5’- ATCGTTGCTGCTGGTGTTG-3’ and 5’-AGGCGATGTGAGCTGTGTG-3’, for N2B isoform; 5’-TATGCCACAACAGCAGGAGA-3’ and 5’-AAGCCTTGCAGGTGTATTCG-3’, for N2BA isoform; 5’-TATGCCACAACAGCAGGAGA-3’ and 5’-TTTGGTCTTTGCAGCCTCTT-3’.

### Overexpression of *TTN* Ig domain

A wild-type (WT) *TTN* cDNA fragment covering A156 (from bp89426 to bp89711 of NM_133378 corresponding to aa29735-aa29829) or *TTN* mutants carrying the A to T change (E29783V mutation equivalent of the Medaka mutation D23186V) was cloned into the pCS2+ expression vector ^31^. A capped RNA linked to the cDNA for GFP was synthesized by a MMESSAGE mMACHINE Kit (Ambion), and injected into the cytoplasm of one cell stage embryos at a final concentration of 500 ng/µl.

### Cloning of titin fragment

We prepared fragments of human *TTN* by PCR from a *TTN* A156 immunoglobulin (Ig) domain construct as templates. A wild-type (WT) *TTN* cDNA fragment covering A156 (from bp89426 to bp89711 of NM_133378 corresponding to aa29735-aa29829) domain was obtained by PCR from cDNAs prepared from total heart RNA, and the A to T substitution mutant (E29783V mutation equivalent of the Medaka mutation Asp23186Val) was created by the primer-mediated mutagenesis method.

### Constructs for epitope-tagged proteins

The cDNA fragments of WT and mutant *TTN* was cloned into 3HA-taggged pcDNA3-NTAP. Full length MURF1 fragment in a Myc-tagged pCMV-Tag3B construct,^32^ was cloned into Flag-tagged pcDNA3-NTAP. The Myc-tagged ubiquitin construct was a kind gift from Dr. Nakada. These constructs were sequenced to ensure that no errors were introduced.

### Cellular transfection, protein extraction and immunoprecipitation (IP)

A total of 8 µg of plasmid DNA was transfected into semi-confluent HEK293FT cells on 6-cm-diamitor culture dishes using Lipofectamine 2000 (Invitrogen). Epoxomicin or DMSO was added to the culture 4 h before lysate preparation. At 24 h after transfection, cells were lysed in 400 µl RIPA buffer (25 mM Tris-HCl pH 7.6, 150 mM NaCl, 1% NP-40, 1% sodium deoxycholate, 0.1% sodium dodecyl sulfate (SDS)) with protease inhibitor, incubated on ice for 30 min and centrifuged at 15000 rpm for 30 min. Supernatants were subjected to western blot (WB) and immunoprecipitation (IP). For IP, lysates were diluted 1:10 using TGN buffer (50 mM Tris-HCl pH7.5, 150 mM NaCl, 1% Tween-20, 0.3% NP40) and precleaned by protein G PLUS-Agarose (Santa Cruz). The precleaned lysates were incubated with 1 µg Anti-HA high-Affinity (3F10, Roche) or normal IgG (GE healthcare) for 3 h and combined with 20 µl protein G PLUS-Agarose (Santa Cruz). The lysates with agarose-bead complexes were incubated overnight with rotation. The bead complexes were washed in TGN 4 times and twice in PBS. The beads were suspended in 2 x Laemmli sample buffer (BioRad) containing 15% 2-mercaptoethanol and boiled for 10 min.

### Western blotting

Cellular lysates and IPs were separated by SDS–polyacrylamide gel electrophoresis (PAGE) and transferred to polyvinylidene fluoride (PVDF) membranes. After a pre-incubation with 3% skim milk in Tris-buffered saline (TBS), the membrane was incubated with primary antibody overnight. After TBS washing, the membrane was incubated with an appropriate secondary antibody (anti-rabbit or anti-mouse IgG HRP-conjugated Ab (1:10,000 in 3% skim milk in TBS, GE Healthcare), mouse TrueBlot ULTRA: Anti-Mouse Ig HRP (1:1000 in 3% skim milk in TBS containing 0.1 % Tween-20 (TBST)) for 1 h. After five more washes in TBST, signals were visualized using the Chemi-Lumi One Super (Nacalai) and Luminescent Image Analyzer LAS-3000 mini (Fujifilm, Tokyo, Japan).

### Antibodies

We used the following primary antibodies: First Antibodies: Flag (1:1000 in 5% BSA/TBST, M2, F7425, Sigma), HA (1:2000 in 3% skim milk/TBST, clone HA-7, Sigma), myc (1:400 in 5% BSA/TBST, clone 9E10, Santa Cruz), GAPDH (1:1000 in 5% BSA/TBST, 14C10, rabbit mAb, Cell Singaling), and the following second antibodies: ECL anti-mouse IgG, peroxidase-linked species-specific F(ab’)2 fragment (from sheep) (1:10,000 in 3% skim milk/TBST, NA9310, GE Healthcare) ECL anti-rabbit IgG, peroxidase-linked species-specific F(ab’)2 fragment (from donkey) (1:10,000 in 3% skim milk/TBST, NA9340, GE Healthcare) Mouse TrueBlot ULTRA: Anti-Mouse Ig HRP (1:1,000 in 3% skim milk/TBST, eBioscience). IPs used the anti-HA high Affinity (3F10, Roche) normal mouse IgG (GE Healthcare)

### Clinical subjects

We studied 96 genetically unrelated Japanese patients with HCM, who carried apparent family history of HCM consistent with an autosomal dominant trait. They were diagnosed with HCM according to the diagnostic criteria as described previously.^33^ They had been analyzed for mutations in the known disease genes for HCM, i.e., genes for contractile elements, Z-disc elements, and I band elements, and no disease-associated mutation was found.^33^ Control subjects were 400 healthy Japanese individuals selected at random. Blood samples were obtained from the subjects after written informed consent and the research protocol was approved by the Ethics Review Committee of Medical Research Institute, Tokyo Medical and Dental University, Japan.

### Mutational analysis

Genomic DNA extracted from blood samples of each individual was subjected to PCR in exon-by-exon manner by using primer-pairs specific to exons 296 to 307 of *TTN* (GenBank Accession No. NM_133378), which encode for titin at the A-band-M-line transition zone including the titin-kinase domain. Sequences of primers and PCR conditions used in this study are listed in Supplementary Table 1. PCR products were analyzed by direct sequencing using Big Dye Terminator chemistry (Applied Biosystems) and an ABI3100 DNA analyzer (Applied Biosystems).

### Co-immunoprecipitation assay

We used cDNA fragments of human *TTN* and MURF1 by PCR from *TTN* A156, A160-161, and A168-170 immunoglobulin (Ig)-like domain constructs and a MURF1 full-length construct, respectively, as templates for PCR. A wild-type (WT) *TTN* cDNA fragment covering A156 (from bp89426 to bp89711 of NM_133378 corresponding to aa29735-aa29829), A160-161 (from bp90626 to bp91202 corresponding to aa30134-aa30326), A168 (from bp93006 to bp93287 corresponding to aa30928-aa31021), A169 (from bp93288 to bp93581 corresponding to aa31022-aa31119), or A168-A170 (from bp93006 to bp93908 corresponding to aa30928-aa31228) domains was obtained by PCR, and *TTN* mutants carrying A to T (E29783V mutation equivalent of the medaka mutation D23186V), T to G (an HCM-associated S30186A mutation), or T to A (another HCM-associated D30994N mutation) substitutions were created by the primer-mediated mutagenesis method. A full-length MURF1 cDNA fragment spanning from bp137 to bp1198 of GenBank Accession No. NM_032588 (corresponding to aa1-aa354) was obtained by PCR. The *TTN* cDNA fragment was cloned into pEGFP-C1 (Clontech, CA, USA), while the MURF1 cDNA fragment was cloned into myc-tagged pCMV-Tag3 (Stratagene, CA, USA). These constructs were sequenced to ensure that no errors were introduced.

Cellular transfection and protein extraction were performed as described previously,^33^ and co-IP assays were done using the Catch and Release v2.0 Reversible Immunoprecipitation System according to the manufacturer’s instructions. IPs were separated by SDS-PAGE and transferred to nitrocellulose membrane. After a pre-incubation with 3% skim milk in PBS, the membrane was incubated with primary rabbit anti-myc polyclonal or mouse anti-GFP monoclonal Ab (1:100, Santa Cruz Biotechnology), and with secondary goat anti-rabbit (for polyclonal Ab) or rabbit anti-mouse (for monoclonal Ab) IgG HRP-conjugated Ab (1:2000, Dako A/S, Glostrup, Denmark). Signals were visualized by Immobilon Western Chemiluminescent HRP Substrate (Millipore, MA, USA) and Luminescent Image Analyzer LAS-3000mini (Fujifilm, Tokyo, Japan), and their densities were quantified using Multi Gauge ver3.0 (Fujifilm, Tokyo, Japan). Numerical data were expressed as means ± SEM. Statistical differences were analyzed using Student’s *t* test for paired values. A *P*-value of less than 0.05 was considered to be statistically significant.

### *In vitro* ubiquitination assay

*In vitro* ubiquitination reactions contained as E1 ligase 100 nM UBE1 (Boston Biochem), and 1µM UbcH5c as E2 ligase (Boston Biochem), 150 µM ubiquitin, 4 mM ATP, and 2,5 µM recombinant titin A168-170 fragment. Reactions were started by addition of 2 µM MURF1 or MURF2, respectively. After incubation for 4 hours at room temperature, samples were boiled in SDS loading buffer, products separated by SDS-PAGE, and blotted onto nitrocellulose membranes. Titin-specific products were detected by using antibodies directed to titin A169, or titin A168-170.

## Non-standard Abbreviations and Acronyms

amhc: atrial myosin heavy chain
BNP: brain natriuretic peptide
cmlc2: cardiac myosin light chain 2
co-IPs: coimmunoprecipitations
DCM: dilated cardiomyopathy
dpf: days post fertilization
ENU: N-ethyl-N-nitrosourea
fli1: friend leukemia integration 1 transcription factor
GFP: green fluorescent protein
HCM: hypertrophic cardiomyopathy
Ig: immunoglobulin
IP: immunoprecipitation
LG: linkage group
MVP: motion vector prediction
MURF: muscle-specific ring finger protein
NRC: neonatal rat cardiomyocytes
nsh: *non-spring heart* Medaka mutant
PCR: polymerase chain reaction
ROI: region of interest
TTN: titin gene
UPS: ubiquitin proteasome
vmhc: ventricular myosin heavy chain
WT: wild-type

## Acknowledgments

We thank the National Bio Resource Project for supplying medaka strains and other support. We thank Dr. T. Hayakawa and T. Kunihiro from Sony Corporation for their technical help in the high-speed video recording of medaka embryos and in the image analysis using MVP method. We are also grateful to Ms. Ono for maintenance of medaka.

## Movie legends

**Movie 1. Contracting heart of a WT embryo.**

Representative frontal view of a WT heart at 5 dpf. Atrium was on this side and ventricle was deeper side. Blood flow was observed in the myocardial wall.

**Movie 2. Contracting heart of a *nsh* mutant embryo.**

Representative frontal view of a mutant heart at 5 dpf. The *nsh* mutant heart had lost elasticity, beat in a rigid manner and showed increased thickening of the myocardial wall, even without blood flow.

**Supplemental Figure I.**
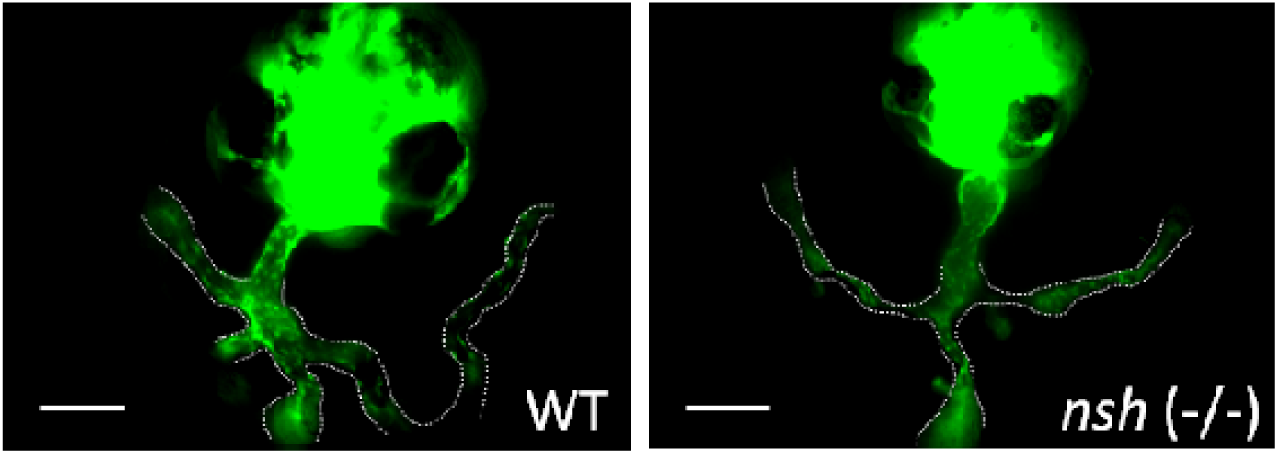
Normal vascular formation was observed in the *nsh* mutant. Crossing with *fli1*-GFP transgenic medaka revealed normal vascular endothelial cells and normal blood vessel formation in the WT and *nsh* mutant at 3 dpf. Scale bars, 200 µm.

**Supplemental Figure II.**
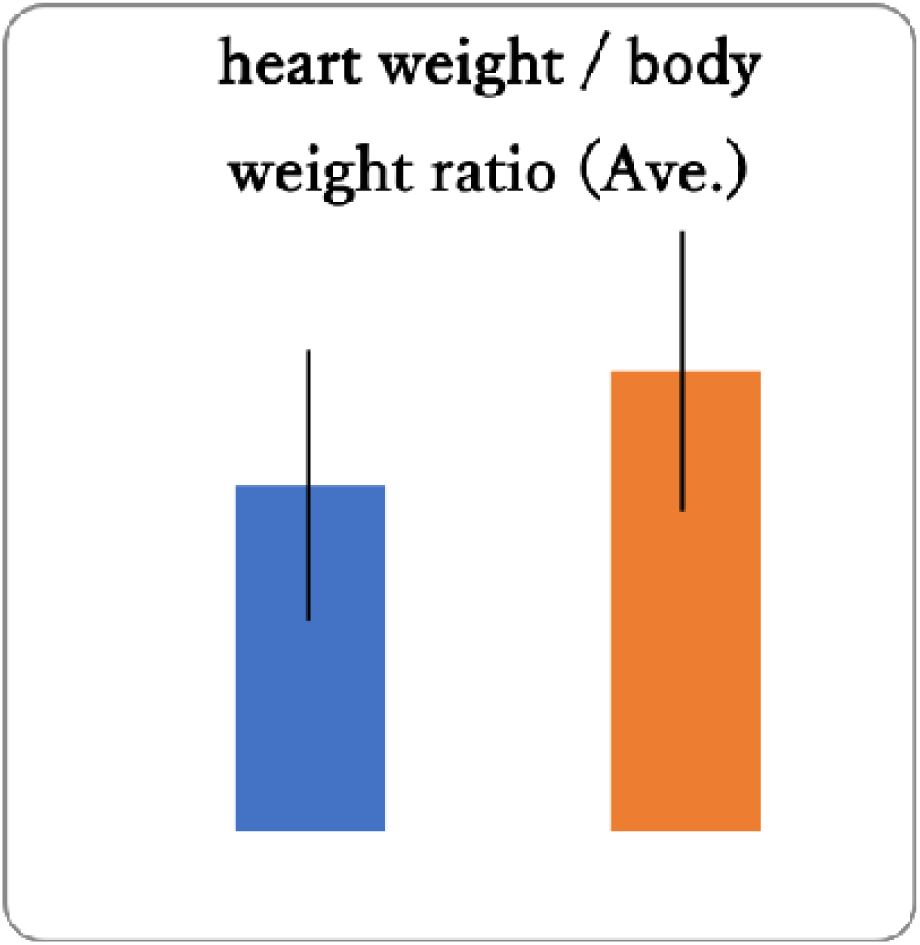
Measurement of heart weight/body weight ratios.

**Supplemental Figure III.**
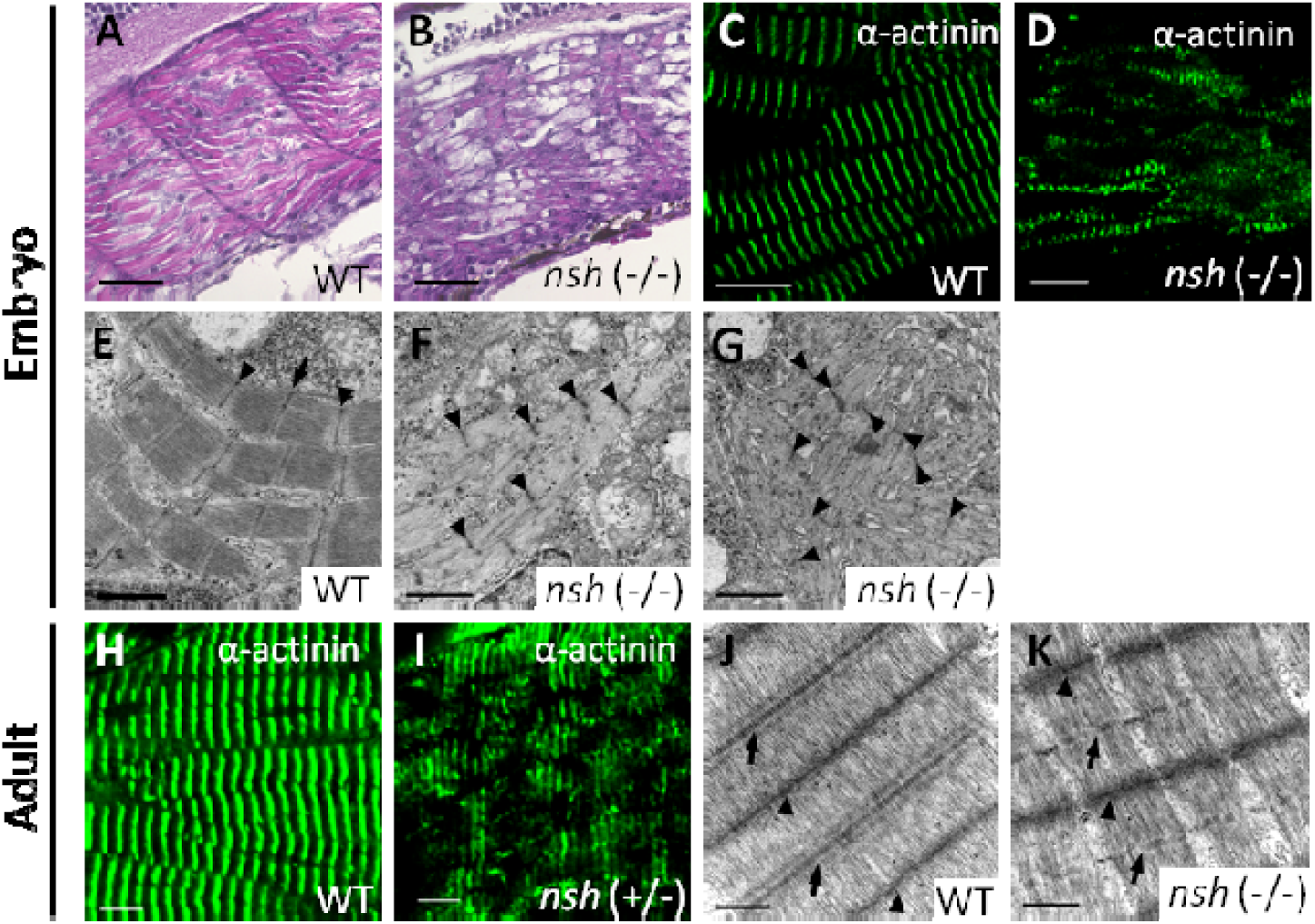
Sarcomeric structures were disrupted in the *nsh* skeletal muscles. A and B, Longitudinal sections of medaka embryonic skeletal muscle at 5 dpf, stained with hematoxylin/eosin. WT muscle showed regularly aligned myofibers, whereas the *nsh* myofibers were disorganized. C and D, Embryonic skeletal muscle of WT and *nsh* mutant (*nsh* −/−) at 3 dpf stained for α-actinin. Sarcomeric structures were disrupted in the *nsh* mutant (D). E through G, Transmission electron microscopy of medaka embryonic skeletal muscles at 5 dpf. (E) In the WT skeletal muscles, myofibrils were assembled regularly. F and G, In the *nsh* mutant, skeletal muscles showed severely disturbed sarcomeric integrity with ruptured myofibrils, blurry Z-discs and M-lines, irregular sarcomere length, lack of thick filaments, and pronounced myocyte disarray (G). H and I, Adult skeletal muscle from WT and *nsh* heterozygotes (*nsh* +/−) stained for α-actinin. Patchy staining was observed occasionally in the *nsh* heterozygotes (I). J and K, Transmission electron microscopy of medaka adult skeletal muscles. J, WT demonstrated well-defined Z-discs and M-lines in skeletal myofibrils. K, Skeletal myofibrils were visible in the *nsh* heterozygotes. Although the mutant displayed well-formed Z-discs and mature-appearing myofibrils, it also demonstrated occasionally disrupted M-lines. Z-discs (arrowhead) and M-lines (arrow) are indicated. Scale bars, 30 µm in (A and B), 10 µm in (C and D), 1 µm in (E through G), 5 µm in (H and I), and 0.5 µm in (J and K).

**Supplemental Figure IV.**
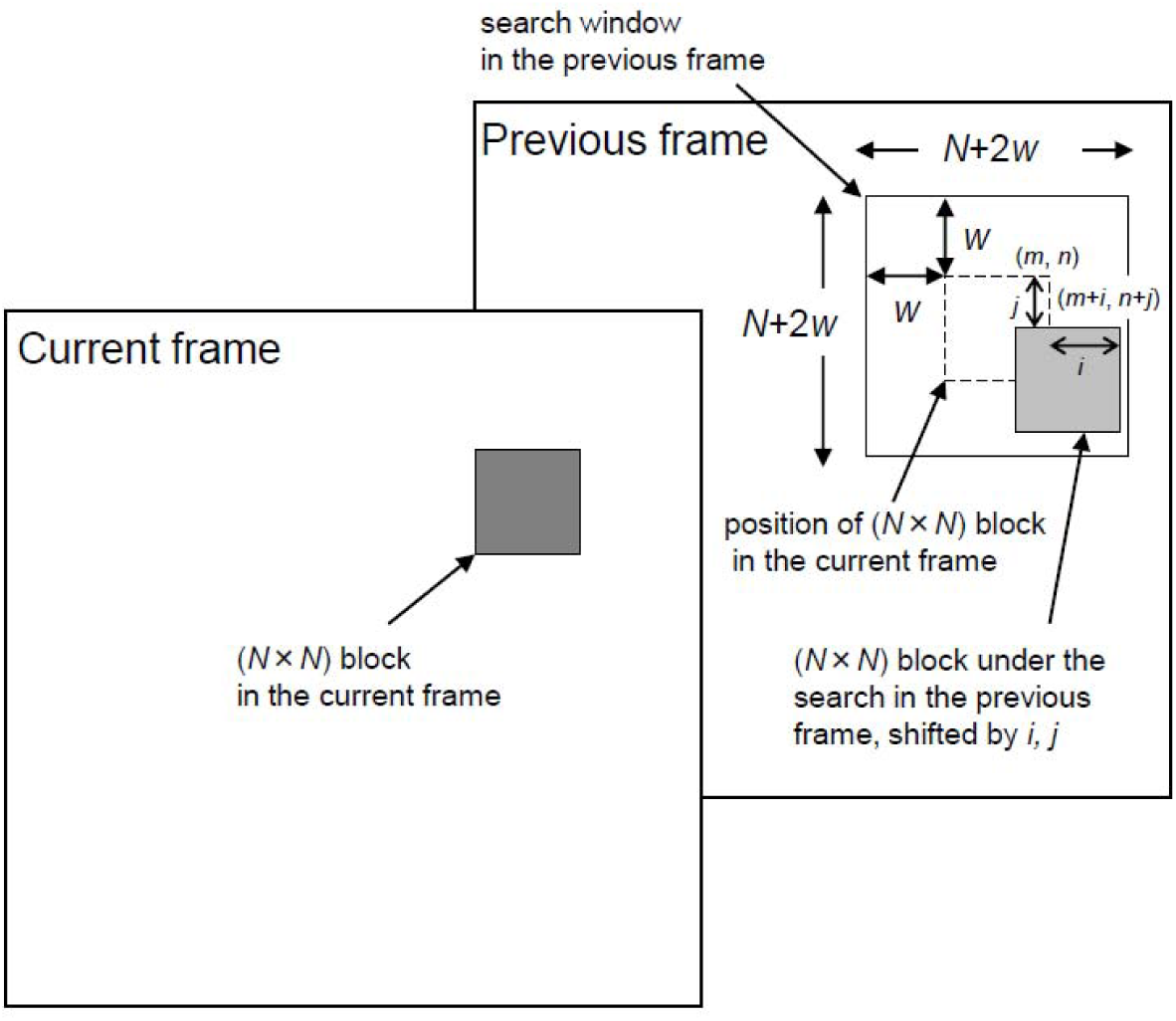
The Schematics of sequential frames in a search window. Each frame was divided into square blocks of *N* × *N* pixels. For a maximum motion displacement of *w* pixels per frame, the current block of pixels was matched to the corresponding block at the same coordinates in the previous frame within a square window of width *N* + 2*w*. In this study, we set the parameters, N = 16 and W = 4.

**Supplemental Table I.**
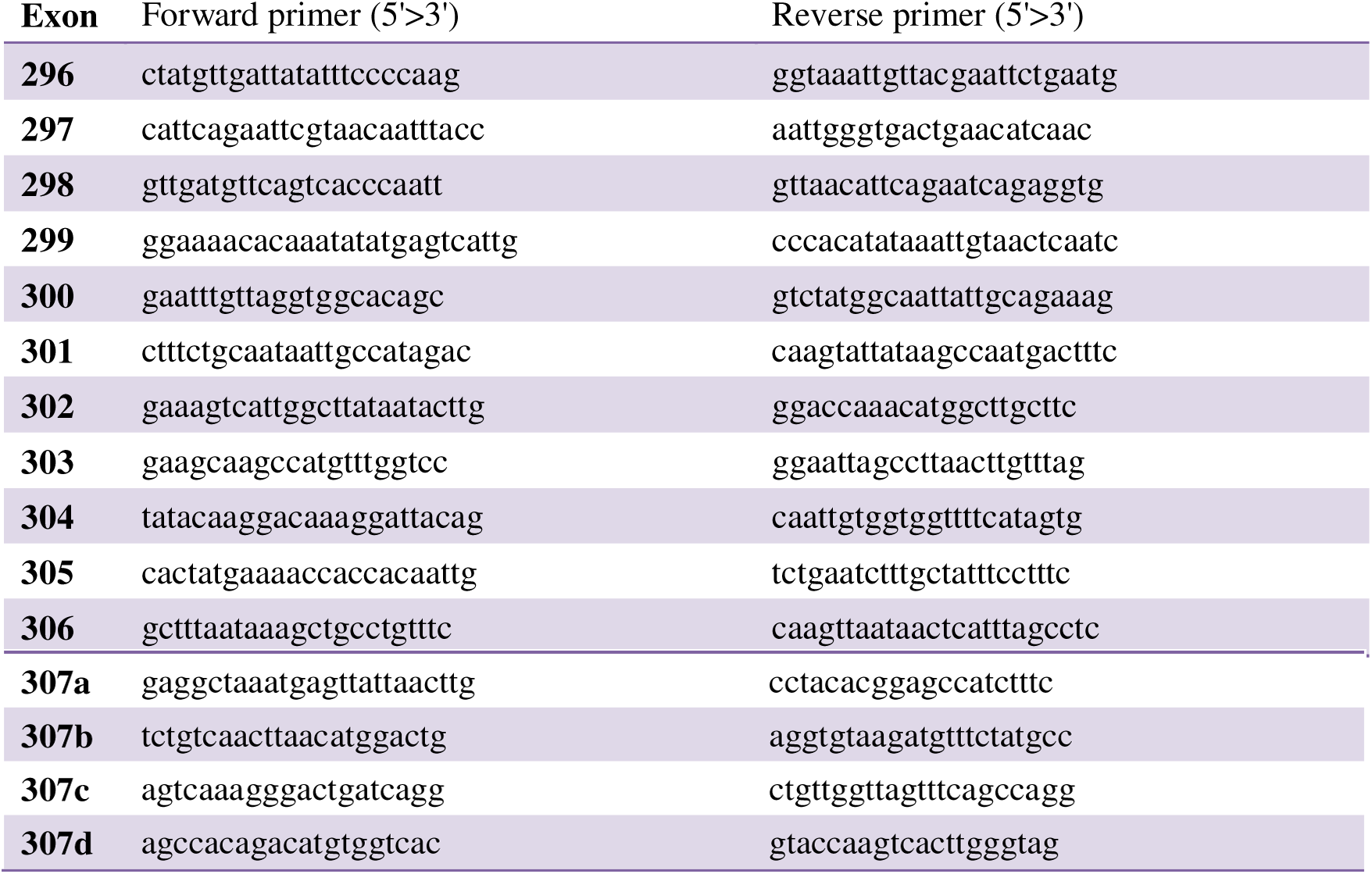
Primers used for PCR amplification and sequencing of human *TTN*.

